# Efficient graphene oxide coating improves cryo-EM sample preparation and data collection from tilted grids

**DOI:** 10.1101/2021.03.08.434344

**Authors:** Avinash Patel, Daniel Toso, Audrey Litvak, Eva Nogales

**Affiliations:** Biophysics Graduate Group, UC Berkeley; Molecular Bioscience Department, Northwestern University; QB3 Institute, UC Berkeley; Molecular and Cell Biology Department, UC Berkeley; MBIB Division, Lawrence Berkeley National Laboratory; HHMI, Berkeley

## Abstract

Recent technical developments have made single particle cryo-EM a major structural biology technique, especially in the characterization of challenging samples that resist crystallization, can only be obtained in small amounts, or suffer from compositional or conformational heterogeneity. However, a number of hurdles that often challenge sample preparation still need to be overcome in order to increase the applicability and throughput of cryo-EM. These technical hurdles include obtaining enough particles per image, with close to random orientation, and without damage from interaction with the air-water interface. While coating EM grids with graphene oxide is a promising procedure for the improvement of sample preparation, it suffers from some technical problems that limit its applicability. We have modified the established drop cast method for adhering graphene oxide onto holey patterned grids to increase graphene coverage. Our method relies on the use of a polycationic polymer to coat the surface of the grid prior to graphene oxide application, thereby preventing the repulsion of the negatively charged graphene oxide sheets from the negatively charged grid surface. With this improved preparation method, we show that graphene oxide supports can increase the number of particles in the field of view by an order of magnitude with respect to open holes, while keeping them away from the damaging air-water interface. We also show how graphene oxide coated gold foil grids can be used to collect tilted cryo-EM data in order to overcome preferred orientation issues, without experiencing the large amount of drift observed with conventional amorphous carbon supports, thus allowing data collection that can lead to high-resolution reconstructions.

## Introduction

Developments in direct electron detector technology and computational image processing tools in the past decade have made single particle cryo-electron microscopy (cryo-EM) a leading method for the determination of macromolecular structures alongside X-ray crystallography and nuclear magnetic resonance spectroscopy. As a single particle technique, cryo-EM has the advantage of being able to determine the structures of large, flexible, and heterogeneous molecules, as each particle can be classified and individual domains refined separately, all with the need for only small amounts of sample ^1^.

Though cryo-EM enables us to look at many new samples, it does come with its own difficulties. A major barrier in obtaining a high-quality reconstruction is preparing a high-quality sample, in which the particles are densely packed, yet separated, randomly oriented, and undamaged ^2^. In order to fulfill these needs, many methods have been devised and used to varying levels of success.

Increased sample concentration is the easiest way to image a greater number of particles without compromising data quality by lowering the imaging magnification or increasing the data collection time at a corresponding increased cost. If concentrated samples are not feasible during biochemical preparation, the main method of obtaining more particles is to use a support layer over the EM grid that binds the particles, effectively concentrating them by adsorption. The simplest and most popular support is a thin layer of amorphous carbon ^3^, but graphene ^4–8^, graphene oxide ^9–11^, or lipid monolayers ^12,13^ have also been used. Adsorption to a support layer generally results in increased particle numbers with increasing incubation times ^14,15^. In addition to a simple support layer, affinity grids have been developed with specific interacting groups to capture molecules in solution ^13,16–19^. In the absence of a continuous support layer, molecules may preferentially localize to the holey support layer of the grid. Another approach to increase particle numbers in open holes has therefore been to passivate the grid to make it easier for the applied sample to enter/remain in the hole ^20–22^.

The damaging effect of the hydrophobic air-water interface results in the breakdown of many fragile complexes, causing subunits to dissociate and/or proteins to denature ^6,23^. There are several approaches that aim to solve this problem, including the addition of small amounts of detergents to the sample ^24^, stabilizing the protein complexes by optimizing the buffer conditions (buffers containing sugars and/or glycerol) ^25,26^, crosslinking the sample ^14^, sequestering the particles away from the air-water interface by binding them to a support layer ^16,27,28^, or reducing the exposure time to the air-water interface by rapid freezing of the sample following blotting ^23,29^.

Reconstruction of a three-dimensional object from two-dimensional projection images requires multiple views of the complex. While in solution, particles are tumbling and assume completely random orientations. However, during cryo-EM sample preparation, particles tend to adhere to available surfaces (support layer or air-water interface), which tends to predominantly occur with specific surfaces of the particles, resulting in preferred orientations (e.g. hydrophobic patches will bind the air-water interface). Efforts to overcome this issue include the addition of detergents ^24,30–32^, glow discharging the grid in the presence of different residual chemical groups ^33–35^, coating the grids with different polymers ^36,37^, deforming the grid ^38^, or tilting the grid during data collection ^39^. Except for tilting the grid, none of these methods is generally applicable, leading to lengthy screening procedures that may not always prove effective ^40^. Unfortunately, while tilting has been shown to be most successful to produce multiple views when samples are prepared using open holes ^39^, this type of sample may suffer from the previous two problems, and requires the use of gold foil grids in order to reduce the beam-induced motion ^41^ observed for tilted specimens. While attempts have been made to use amorphous carbon coated grids to collect tilted data, the resulting structures ^42,43^ have not been able to reach the resolution of structures collected in open holes.

Here we show how pretreatment of the grid with a polycationic polymer, such as polyethyleneimine (PEI), can significantly increase the coverage of graphene oxide onto holey patterned grids. We show that such grids can increase particle concentration by an order of magnitude compared to the same sample prepared on open hole grids due to the adsorption of particles onto the graphene oxide sheet, while at the same time being kept away from the damaging air-water interface. Lastly, we demonstrate that graphene oxide-coated gold foil grids can be used to collect tilted cryo-EM data without experiencing large amounts of drift that would otherwise dampen the high-resolution signal.

### Preparation of graphene oxide coated grids

Graphene oxide is a near-ideal support layer for cryo-EM. It produces minimal background in cryo-EM images, is sufficiently hydrophilic to bind biological macromolecules, can be effectively wetted, and blots uniformly during sample preparation ^9^. So far, two main methods have been described to adhere graphene oxide onto the surface of holey carbon grids: the drop cast method ^9,10^ and the surface assembly method ^11^. In the drop cast method, a suspension of graphene oxide is directly applied to a glow-discharged grid, allowing it to adhere to the surface before being washed with water and dried. In our hands, this method leads to limited coating of the gird, making it impractical for automated data collection. The surface assembly method works by forming a monolayer film of graphene oxide on the surface of the water, then lowering the water level until the monolayer coats the submerged grids. While this method is effective in coating the grids, the process can be time consuming and requires careful testing of the amount of graphene oxide needed to form a monolayer.

We have modified the drop cast method to increase the efficiency of graphene oxide adsorption onto the grid surface by coating the grid with polyethyleneimine (PEI) prior to graphene oxide application. This process involves three main steps – grid cleaning, PEI treatment, and GO coating – and it takes approximately 30 minutes to make a batch of grids (∼4-12) (Fig. 1A). Since most commercially available grids come with some amount of plastic or other organic contamination, and are relatively hydrophobic out of the box, the surface of the grid must first be cleaned and rendered hydrophilic. To remove any plastic contamination, the surface of the grids is cleaned using an organic solvent (e.g. chloroform). To render the surface of the grids hydrophilic, they are plasma cleaned in a chamber contain low pressure atmospheric air (Fig. 1 – figure supplement 1A). The plasma treatment ionizes oxygen in the air, which can then oxidize the surface of the grid, rendering it hydrophilic with a slight negative charge ^44–48^. In the second step, the surface of the grid is treated with a buffer solution of PEI, a polycationic polymer. This step allows changing the surface character of the grid so that it has a positive charge capable of attracting negatively charged graphene oxide sheets. We speculate that the reason why graphene oxide does not effectively bind the grid surface without this step is that the two surfaces (plasma cleaned grid and graphene oxide) repel each another as both are negatively charged. In the third and final step, a dilute suspension of graphene oxide sheets is used to coat the surface of the grid.

**Figure 1.**
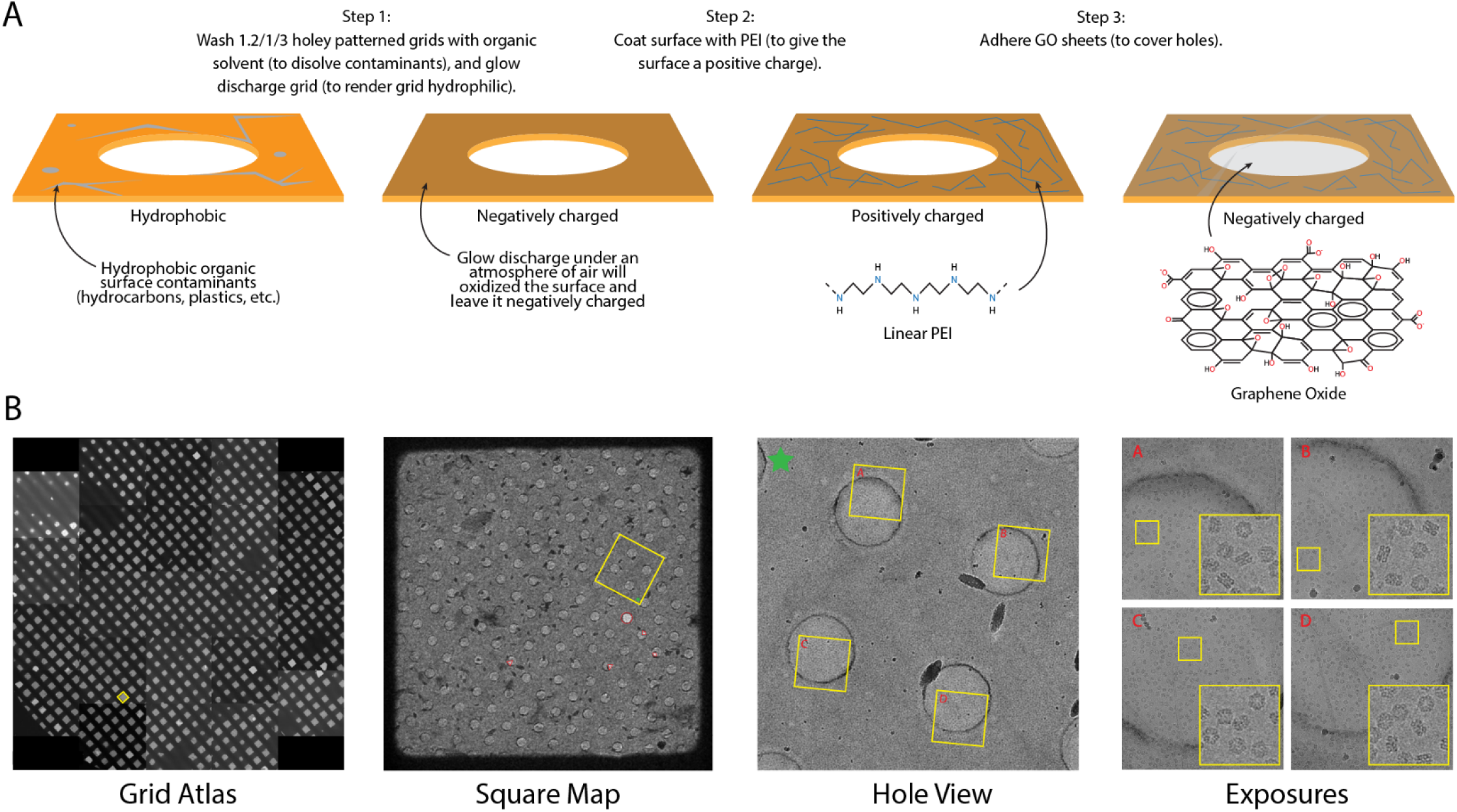
Modified graphene oxide coating method. A. Holey patterned grid preparation and coating workflow. The grids are first washed with organic solvent (i.e., chloroform) and then glow discharged in a chamber contain low pressure atmospheric air. The grids are then coated with a polycationic polymer (i.e., PEI). Finally, the grids are coated with GO sheets to cover the holes. B. Example of a GO coated grid at several magnifications. The yellow squares indicate the region magnified in the image to the left, or the inset for the right-most panel.

By coating the glow discharged grid with a layer of a positively charged polymer, like PEI, we were able to significantly increase graphene oxide coating (Fig. 1B). We typically use regularly patterned grids made with a carbon or gold film. We find that we get near complete coating with smaller hole patterns (2/2 or smaller), but larger sizes can also be used. We also find that the most critical part for the success of this protocol is having a good batch of graphene oxide solution. (Fig. 1 – figure supplement 1B).

### Particle adsorption of various specimen types

In order to compare the sample properties of graphene oxide (GO) coated grids with open-hole (OH) or amorphous carbon (AmC) coated grids, we prepared cryo-EM samples of erythrocruorin from *Eisenia fetida* using suitably pre-treated 1.2/1.3 UltrAuFoil grids. We found that a two-minute incubation time of the sample on the GO and the AmC grids, using 0.08 mg/mL erythrocruorin (22nM), resulted in densely packed particles on the grid, with slightly higher density on GO than AmC (Fig 2A-B, Fig. 3A). The open hole grid, on the other hand, required .8 mg/mL erythrocruorin (220nM) to achieve roughly a quarter of the particle density seen for GO coated grids under otherwise identical conditions (Fig 2C, Fig. 3A). The need for significantly less sample for GO (and AmC) versus OH grids has been observed previously ^11^ and is likely a result of particles adsorbing to the GO and the AmC support films.

**Figure 2.**
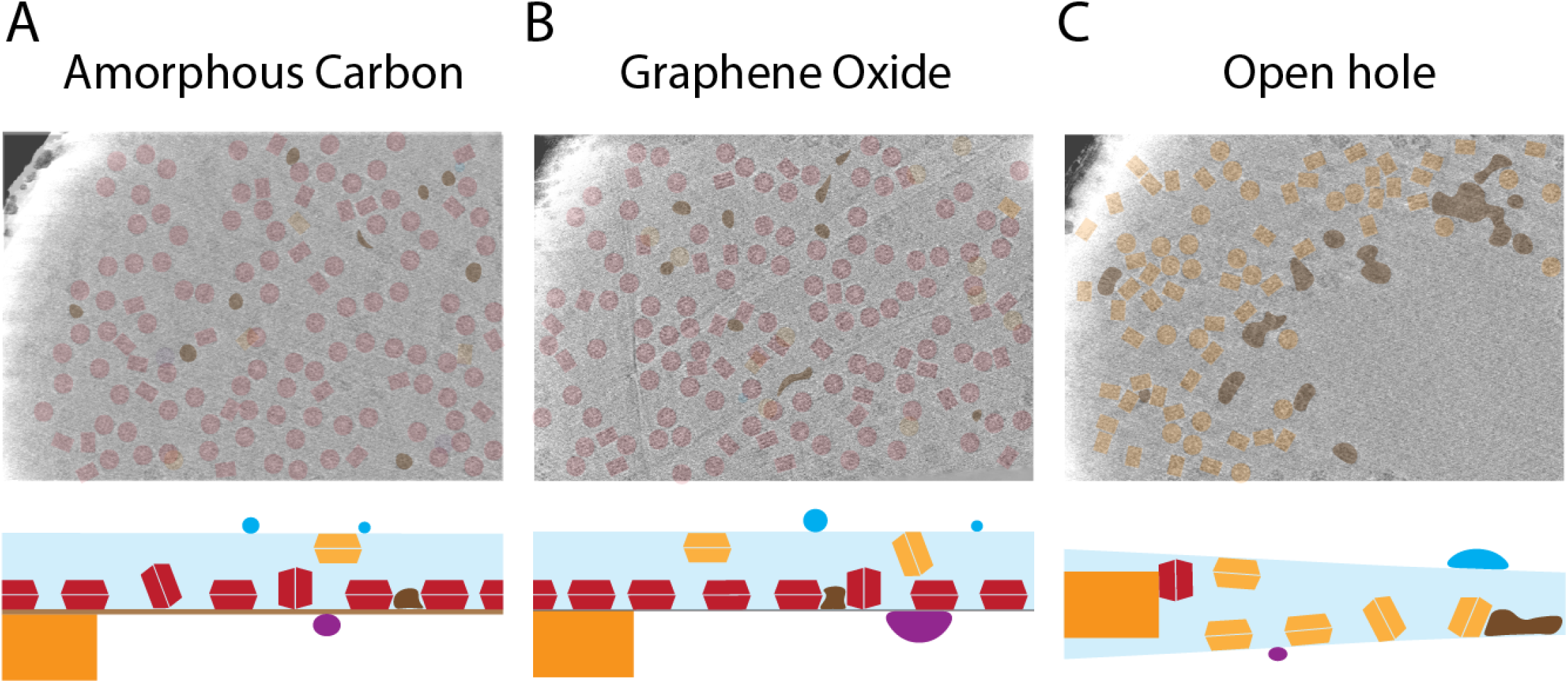
Particle positions on different grid types. Projections in the Z-direction of tomographic reconstructions with segmented particles for samples prepared on **(A)** amorphous carbon, **(B)** graphene oxide and **(C)** open hole grids (top), with cartoon representations of the grids that illustrate how the particles were distributed in the vitreous ice layer (bottom). Particles are colored based on their distance from the lower surface: red - particles bound to support layer, orange - particles bound to air-water-interface, blue/purple – crystalline ice on top or bottom of the vitreous ice layer, brown – damaged and aggregated complexes.

**Figure 3.**
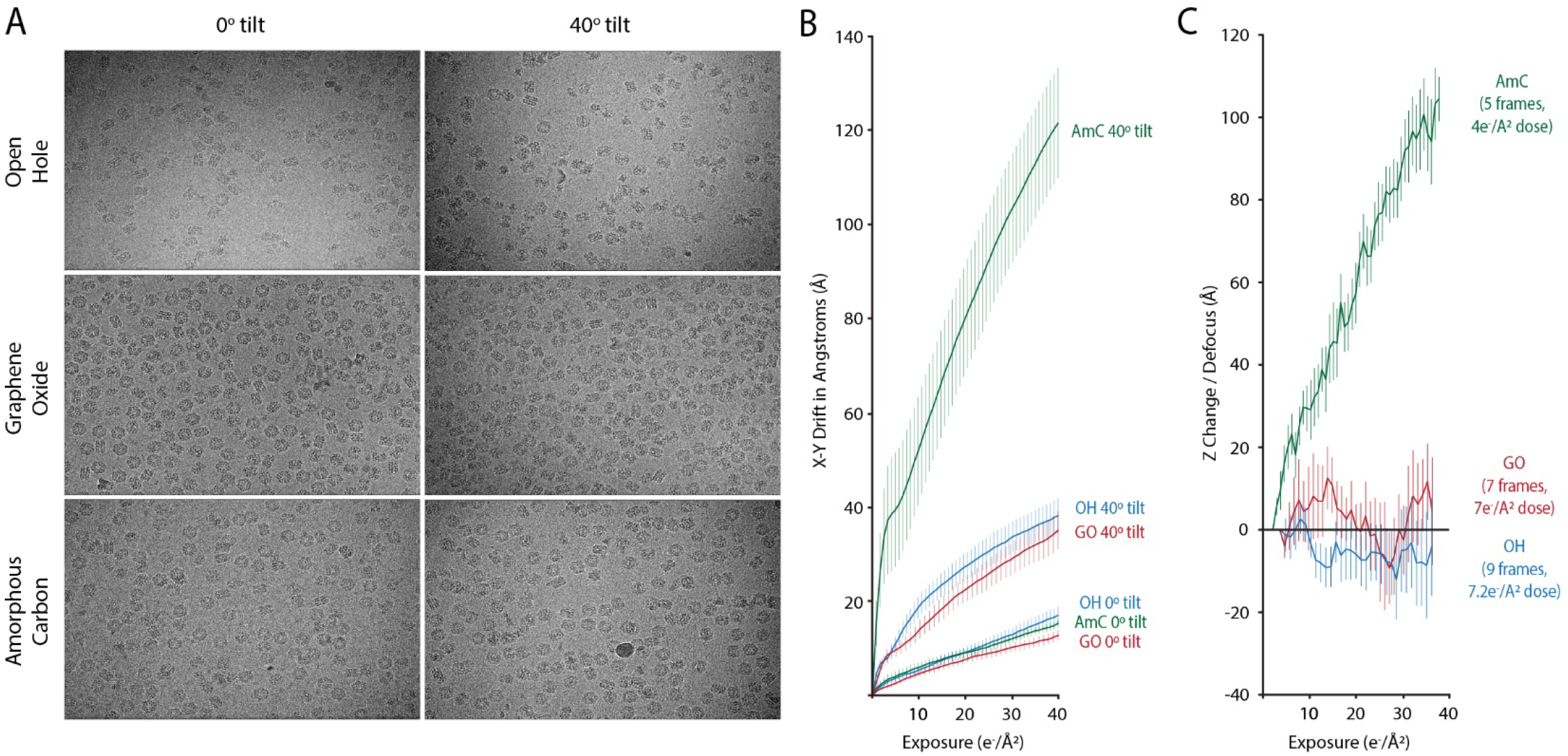
Sample drift during tilted data collection. A. Representative micrographs at 0° and 40° stage tilt for OH, GO, and AmC gold foil grids. B. Tracks represent the average drift for image collected on OH, GO, and AmC grids, at both 0° and 40°. Tracks are averages from ten images, with the vertical lines representing the standard deviation. C. Measured change in defocus during exposure. For the collected movies, a rolling average of aligned and averaged frames was used to increase the signal of the Thon rings for CTF estimation. Tracks are averages from ten images, with the vertical lines representing the standard deviation.

To ascertain whether the increased number of particles on GO and AmC coated grids is due to adsorption of particles onto the support layer, we collected tomograms for each of the three samples. For the GO and AmC coated grids, most of the particles were bound to the support layer, with the remaining particles bound to the air-water interface (Fig 2A; Fig. 2 – figure supplement 1). In stark contrast to the GO and AmC coated grids, the OH grid had all its particles bound to the two air-water interfaces, with a majority of the particles bound to the side away from the blotting paper (Fig 2C; Fig. 2 – figure supplement 1).

The use of support films can significantly increase the particle concentration on the grid, and if GO is used, the background signal added to the images and the resulting contrast loss will be minimal. These properties are particularly useful when the amount of sample is insufficient for the preparation of open-hole grids. An added advantage is that the hydrophilic support can also help to prevent sample damage by constraining particles to the support film and away from the damaging hydrophobic air-water-interface.

### Grid tilting with specimen support layers

For many samples, preferential orientation is a major barrier to obtaining a three-dimensional cryo-EM reconstruction. Even when a reconstruction can be generated, its quality may greatly suffer from anisotropy due to a limited number of views ^39,49,50^. Of the various approaches used to increase angular distribution for a cryo-EM sample, tilting is the only one guaranteed to give additional views. However, to achieve high resolution the collection of tilted specimens has been shown to be effective only when the sample is prepared in open holes using gold foil grids that minimize beam-induced motion ^39^. To test the effect of having a support layer over the gold foil grid, we collected images at 0° and 40° tilts from OH, GO, and AmC grids (Fig. 3A).

To determine the amount of in-plane (XY) drift, images were collected with a total exposure of 40e-/Å^2^, fractionated into 40 or 50 frames that were then aligned to measure the amount of drift. Ten images were collected for each condition (AmC, GO, and OH) at both 0° and 40° tilt. The drift between every consecutive pair of frames was then averaged, and the standard deviation plotted (Fig. 3B). For all the images collected at 0° tilt, the XY drift was measured to be ∼15Å (12.8Å for GO, 15.3Å for AmC, and 17Å for OH). For the 40° tilt images, the average drift was approximately double that of the 0° tilt images for the GO (35.3Å) and OH (38.3Å) grids, but was ∼8 times higher for the AmC (120.7Å) grid.

To determine the potential cause for the increased drift for the AmC grids, we measured the change in defocus over the exposure for the 0° tilt images (Fig. 3C). This was meant to measure the amount of the sample motion in the Z direction during the exposure. We found that in images with GO and OH, there was some amount of fluctuation in the defocus (∼±10Å), but the defocus did not change with dose, while the AmC images increased in defocus by ∼100Å over a total dose of 40e-/Å^2^ exposure. Thus, the large increase in drift for the tilted AmC coated grids can in part be explained by Z motion as reflected in the increase in the defocus seen for these grids. However, given that the change in defocus is ∼100Å and the collected images were tilted 40°, the contribution to the XY drift would only be ∼60Å. The change in defocus also does not explain the increased drift observed in the tilted images for OH and GO images, as there is little change in defocus for these samples.

The expected resolution limitation caused by sample drift can be approximated to be equal to the amount of inter-frame drift. Hence, if 1Å drift occurred during the capture of a single frame, the resolution limit within that frame is limited to 1Å in the direction of the drift. Therefore, in the case of the AmC grid collected at 40° tilt, the theoretical resolution limit would be 3Å, given the used collection settings (1e-/Å^2^/frame). However, this is likely optimistic as there is an even greater amount of drift in the earlier frames, when most of the high-resolution information in the sample is yet been reduced due to radiation damage ^51–53^. To compensate for the increased drift, more data can be collected, or images can be collected at a faster frame rate so that the drift per frame is reduced. However, the effectiveness of using faster frame rates may be limited as the ability to align images may be reduced due to the consequent lower signal-to-noise ration of the individual frames.

### Reconstructions from GO-Au grids

To test the effectiveness of tilted data collection using GO coated UltrAuFoil grids, we collected a dataset of erythrocruorin at 40° tilt on a Titan Krios equipped with a K2 Summit direct electron detector mounted behind an energy filter, as described by Tan et al. (Fig. 4) ^39^. Initial processing using per-micrograph CTF estimation resulted in a 4.9Å resolution reconstruction (defocus gradient ∼3000Å). Correcting the CTF on a per-particle basis improved the resolution to 3.6Å. While the improvement was significant, it should be noted that per-particle CTF estimation does not need to be determined from the start, as this may not be very accurate for some smaller samples. After aberration correction, classification, and polishing, the final resolution obtained was 2.9Å.

**Figure 4.**
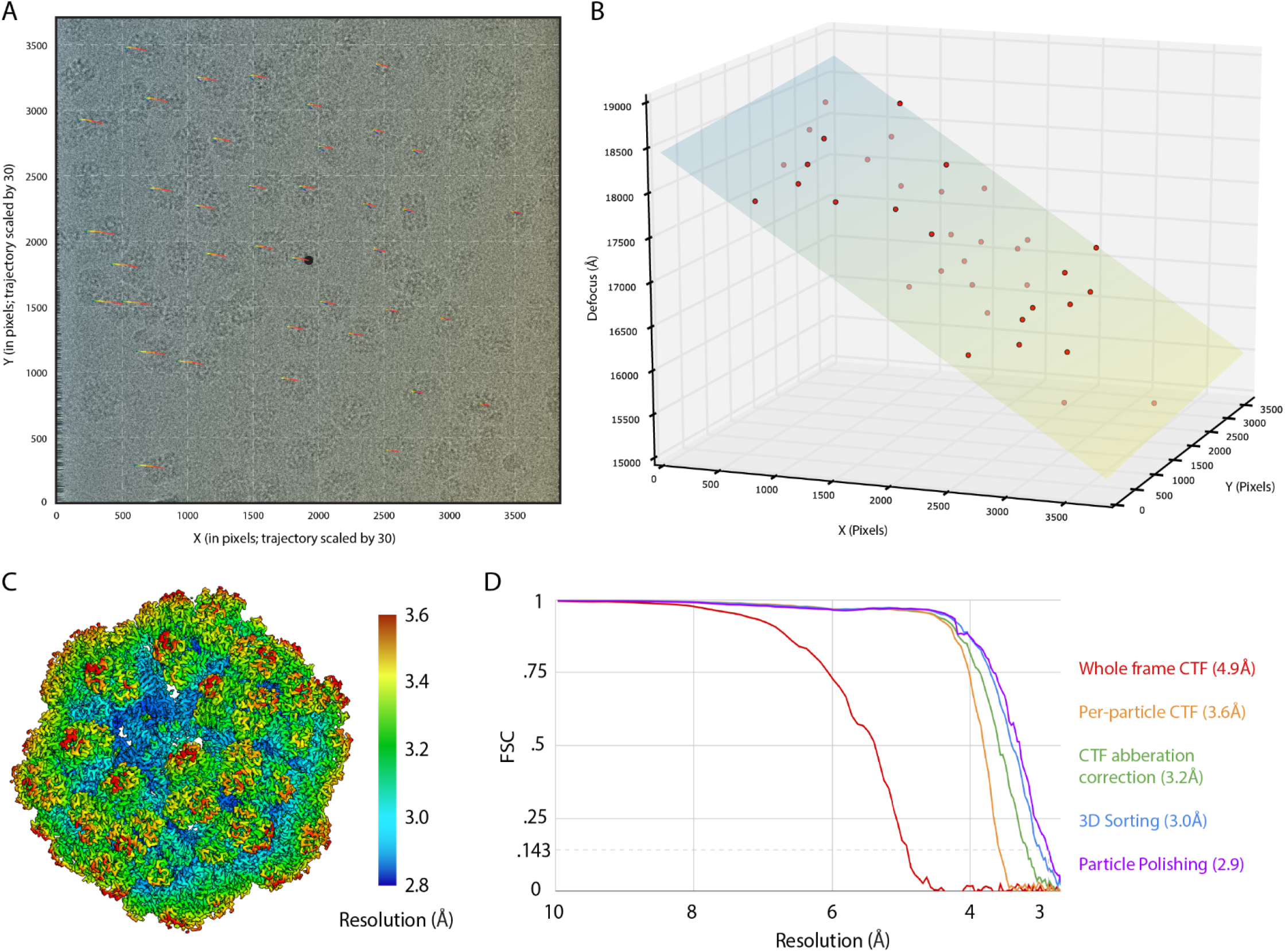
Collection of titled data on GO coated gold foil grids. A. Representative erythrocrurin micrograph marking the particles that went into the final reconstruction. Color gradient represent the fitted defocus across the grid (same as in Fig. 4B). Colored lines starting from the center of particles represent a 30-fold scaled movement of the particle over the course of the movie. B. Plot of measured defocus across the micrograph with a plane fitted through the points. C. Reconstruction of erythrocruorin colored by local resolution as computed in Relion ^62^. D. Gold standard FSC curves of reconstruction at various stages of data processing.

The modified GO coating method has been used with several other samples of varying size, and typically resulted in reconstructions with better than 3Å resolution ^54^(*to be published*). However, it should be noted that in these other cases, tilting was not required. To demonstrate the capabilities of these GO grids, three other samples were collected: 80S yeast ribosome, human apoferritin and the TAF2 subunit of the human transcription factor IID (Fig. 4 – figure supplement 1). A medium sized dataset of 80S ribosome, a large (3.2MDa) asymmetric complex, was collected and reconstructed to 2.7Å resolution globally local resolutions of 2.6Å for the 60S, 2.9Å for the 40S body, and 3.0Å for the 40S head. A small dataset (266 micrographs) of apoferritin, a medium-sized (470kDa) rigid and the highly symmetric (octahedral) complex was collected and reconstructed to 2.6Å resolution. A large dataset (3413 micrographs) of TAF2 (1-985), a small (110kDa) asymmetric protein, was collected and reconstructed to 2.8Å resolution.

## Conclusion

In summary, our improved GO grid coating method can be used to rapidly and reliably prepare cryo-EM samples, thus facilitating sample preparation that has the potential to overcome traditional hurdles in cryo-EM studies. Of added value, our method does not require any specialized equipment that would not be found in any typical EM facility. We currently prepare a batch of GO coated grids right before freezing samples, only adding several minutes to normal OH sample preparation procedures and significantly reducing the required time needed for making coated grids. While we only show three examples of macromolecules solved using these grids, TAF2 represents a typically challenging cryo-EM sample that is small, asymmetric, and dilute. In addition, we have used this procedure to solve larger, more flexible complexes that we were able to refine to high resolution due to our ability to locally refine smaller, flexible regions (∼100kDa) (*to be published*). Finally, we find that GO-coated gold foil grids can be used to collect tilted data with a drift rate comparable to OH grids and significantly less than AmC coated grids. In conclusion, we foresee that our method can help those working with challenging samples that face issues of low-concentration samples, small or flexible complexes, or may be experiencing orientation bias.

## Methods

### Protein purification

*Saccharomyces cerevisiae* ribosomes were purified from the CBP flowthrough of a TAP tag purification ^55^. The CBP flowthrough was subjected to a 2 mL 10-40% glycerol gradient (20mM HEPES pH 7.9, 150mM KCl, 2mM MgCl) centrifuged for 16 hours at 35k rpm at 4 °C (Beckman TLS-55). 80S ribosome was found to have formed a pellet at the bottom of the tube and was resuspended in 20mM HEPES pH 7.9, 150mM KCl, 2mM MgCl, 10% glycerol, aliquoted, flash-frozen in liquid nitrogen, and stored at −80 °C.

*Homo sapiens* ferritin (Q8TD27) was recombinantly expressed with an N-terminal GST tag in *Escherichia coli*. The codon-optimized human ferritin gene was cloned into pET His6-GST TEV LIC Vector (1G) (MacroLab) ^56^, transformed into BL21 competent cells, and plated on LB-agar plates containing kanamycin (50 μg/mL). The plates were incubated overnight at 37°C. A single colony was transferred to a 50 mL starter culture of Luria-Bertani (LB) broth containing kanamycin (50 μg/mL) and grown overnight at 37 °C with shaking (220 rpm), then used to inoculate two 1 L flasks of LB with kanamycin (50 μg/mL). Cultures were grown at 37 °C with shaking (140 rpm) until the optical density at 600 nm reached 0.4-0.6. The cells were then cooled to 18 °C, and 0.1 mM IPTG was added to induce protein expression. After 12-18 h of incubation at 18 °C, the cells were harvested by centrifugation at 4k rpm for 20 min. The cell pellets were resuspended in PBS and centrifuged at 4k rpm for 20 min, then the cell pellets were flash-frozen in liquid nitrogen and stored at −80 °C. The frozen cell pellets were thawed on ice and resuspended in lysis buffer (50mM HEPES 7.9, 300mM NaCl, 1mM MgCl, 5% glycerol, one protease inhibitor tablet). The cell suspension was lysed using sonication 10 s ON / 50 s OFF for a total processing time of 3 min at power level 60% (QSONICA) and subsequently incubated with benzonase for 10 min at 4 °C with slow rocking. Cellular debris was removed by centrifugation at 18k rpm for 60 min at 4 °C (Beckman JA-20). The lysate was incubated with 5mL of washed and packed GST resin (GE Healthcare) for 1 hour at 4 °C with gentle rocking. The resin was poured into a fritted column and the flow-through fraction was collected. The resin was washed with 50mL of lysis buffer, then eluted with lysis buffer plus 10mM glutathione. The elutions were analyzed using SDS-PAGE, and the fractions containing the protein of interest were combined and concentrated to 2mL at a final concentration of 3.85 mg/mL by repeated centrifugation in an Amicon Ultra Centrifugal Filter at 4k rpm for 5 min at 4 °C (Beckman SX4250). The concentrated protein was incubated overnight with TEV (1:50 mass ratio) with slow rocking at 4 °C to cleave the N-terminal His6-tag. Following the overnight TEV digest, the combined protein elutions were further concentrated to 0.5mL at the same conditions, and SDS-PAGE confirmed the completion of the digest. Potential precipitates were removed from the concentrated protein by centrifugation at 14.2k rpm for 5 min at 4 °C. The digested protein was then loaded onto a Superose 6 10/300 column which was pre-run with the wash buffer (20mM HEPES 7.9, 150mM NaCl). The elutions were analyzed using SDS-PAGE, and the elutions containing the protein of interest were combined, flash-frozen in liquid nitrogen, and stored at −80 °C.

*Homo sapiens* TAF2 (1-985) was recombinantly expressed with an N-terminal MBP tag in *Trichoplusia ni* High Five cells. The codon optimized human TAF2 gene was cloned into pFastBac His_6_-MBP-Asn10-TEV vector (438C) (MacroLab) ^56^ and transformed into DH10 MultiBac competent cells to generate bacmids for baculovirus production. Purified bacmids were transfected into *Spodoptera frugiperda* Sf9 cells using FuGene transfection reagent and amplified for a further two rounds before being used to infect 500mL cultures of High Five cells for protein production. High Five cells were grown for 60 hours post-infection at 28□°C, were then harvested by centrifugation, washed in PBS, flashed frozen, and stored at -80 °C. Pellets were resuspended in lysis buffer (50 mM HEPES pH 7.9, 500□mM NaCl, 2 mM MgCl2, 10% glycerol and lysed by sonication (10s ON / 50s OFF for a total processing time of 3 min at power level 50%) and subsequently incubated with benzonase for 10 min at 4 °C with slow rocking. Cellular debris was removed by centrifugation at 18k rpm for 60 min at 4 °C. Clarified lysate was then passed over 2mL of packed Ni-NTA resin and washed with lysis buffer before being eluted with lysis buffer containing 250mM imidazole. Elutions were pooled and incubated with 1mL of amylose resin overnight. Amylose resin was subsequently washed with lysis buffer and eluted with lysis buffer with 20mM maltose. The elution was then TEV digested overnight, and the digest was incubated with .25mL of Ni-NTA resin. The flow-through of the Ni-NTA resin containing TAF2 was loaded onto an Superdex 200 3.2/300 and run using lysis buffer. Fractions containing pure TAF2 were pooled and concentrated to 4 µM, flash-frozen in liquid nitrogen and stored at −80 °C.

### Graphene oxide grid preparation

Graphene oxide coating was performed on 1.2/1.3 holey patterned carbon or gold film grids (Protochips or Quantifoil). To wash them, grids were placed face-up on a piece of filter paper in a glass dish. A drop of chloroform was added to the top of each grid and allowed to dry; this was repeated once more. The grids were then placed in a small glass petri dish lined with a stainless steel mesh and glow discharged in either a Solarus plasma cleaner (10s, 5W, 80mTorr), Cressington Sputter coater (30s, 15mA, ∼.04 mBar), or Tergeo plasma cleaner (60s, 15W, 210mTorr) in a chamber filled with residual air. The grids were then picked up using an anti-capillary tweezer, and 4uL of 1mg/mL PEI (Polyethylenimine HCl MAX Linear 40K from Polysciences Catalog No. 24765-1 – the higher the molecular weight of the polymer, the better it stays adhered to the grid during subsequent wash steps) buffered in 25mM HEPES pH 7.0 (the buffer was prepared fresh before being applied to the grids by mixing 4uL 10mg/mL PEI with 36uL 27.78mM HEPES pH 7.9) was applied to the top of the grid. The drop was incubated on the grid for 2 minutes and blotted off with a filter paper, and washed twice by adding a 4uL of water on top of the grid and blotting off (first water wash was added to the top of the grid as soon as possible to avoid letting the PEI drop residue from drying on the grid). The grid was returned to a glass petri dish lined with filter paper to dry (at least 2 minutes). The grids were then picked up using anti-capillary tweezers, and a 4uL drop of .2mg/mL GO (Graphene Oxide dispersion at 2mg/mL Sigma-Aldrich 763705-25ML; the suspension aggregates with time, we typically replace our stock every ∼1-3 years when we start observing clumps in the suspension) diluted in water and clarified by centrifugation at 1500g for 1 minute was applied to the top of the grid. The drop was incubated on the grid for 2 minutes and blotted away with a filter paper (we slide the filter while blotting to prevent clogging the filter paper, then continue to the next water wash as fast as possible to avoid the excess GO flakes from drying on the grid surface). We wash twice by adding a 4uL of water on top of the grid and then blot the water off. The grid is then returned to a glass petri dish lined with filter paper and let dry (at least 2 minutes) before using. Grids were typically used an hour after being prepared and always used the same day, as grids tend to become hydrophobic over time. If GO over coverage is observed, the power of the glow discharge should be reduced or additional water washes after GO application performed; conversely, if coverage is insufficient, glow discharge power should be increased or a lower GO dilution should be used for grid application.

### Cryo-EM sample preparation

For open hole cryo-EM sample preparation, grids were first washed with chloroform, dried, and glow discharged in a Solarus plasma cleaner for 20s at 5W and 80mTorr. For sample freezing, 4uL of erythrocruorin at 0.8mg/mL was added to the grid at 4 °C under 100% humidity in a Mark IV Vitrobot (FEI), blotted with a Whatman #1 for 3 sec at 5N force and then immediately plunged into liquid ethane cooled by liquid nitrogen.

To prepare AmC coated grids, grids were washed with chloroform and then coated with a layer of thin carbon ∼2nm thick. Grids were then glow discharged in a Solarus plasma cleaner for 10s at 5W and 80mTorr. For sample freezing, 4uL of erythrocruorin as 0.08mg/mL was added to the grid and incubated for 2 minutes at 4 °C under 100% humidity in a Mark IV Vitrobot (FEI), blotted with a Whatman #1 for 2.5 sec at 0N force and then immediately plunged into liquid ethane cooled by liquid nitrogen.

GO coated grids were prepared as described in the section above, and the freezing conditions were the same as AmC coated grids. For 80S ribosome (unknown concentration) the sample was incubated for 5 minutes, for apoferritin (0.1mg/mL; 210nM) the sample was incubated for 1 minute and for TAF2 (0.22mg/mL; 2uM) the sample was incubated for 3 minutes.

### Cryo-EM data collection

For Fig 1, grids were transferred to a 626 Cryo-Transfer Holder (Gatan) and loaded into an F20 electron microscope (FEI) operated at 120 keV and equipped with a Gatan Ultrascan 4k x 4k CCD camera, and data was collected with Leginon ^57^. High magnification images were collected with a pixel size of 2.29Å/pix.

For Figs 2 and 3, grids were loaded into a Talos Arctica electron microscope (Thermo Fisher Scientific) operated at 200 keV acceleration voltage and equipped with a K3 direct electron detector (Gatan) operated in super-resolution mode. Data were collected with SerialEM ^58^. For tomograms, a bidirectional tilt scheme was used starting at -20° to +59° and then -23° to -57° with a 3° angular step size. Each tilt was collected at -3µm defocus at a pixel size of 1.12Å/pix and a total dose of 3e-/Å^2^. For Fig 3, images were collected at 0° and 40° with a pixel size of 1.12Å/pix for a total dose of 40e-/Å^2^. For AmC and OH grids, movies were collected with 50 frames, and for the GO grid, the movies were collected with 40 frames.

For Fig 4, erythrocruorin grids were loaded into a Titan Krios electron microscope (FEI) operated at 300 keV and equipped with a K2 Summit direct electron detector mounted behind an energy filter (Gatan) operated in super-resolution mode and data was collected with SerialEM ^58^. Images were collected at 40° tilt at a pixel size of 0.90Å/pix with a total dose of 40e-/Å^2^. 80S ribosome, apoferritin and TAF2 grids were loaded into a Talos Arctica electron microscope (FEI) operated at 200 keV and equipped with a K3 direct electron detector (Gatan) operated in super-resolution mode and data was collected with SerialEM ^58^. For 80S ribosome images were collected at a pixel size of 1.12Å/pix with a total dose of 40e-/Å^2^ and for apoferritin and TAF2 images were collected at a pixel size of .69Å/pix with a total dose of 40e-/Å^2^.

### Cryo-EM data processing

The tomographic reconstructions shown in Fig 2 were performed in IMOD ^59^ using the automated patch tracking procedure. Tomographic volumes were reconstructed using five iterations of SIRT and binned 8-fold for visualization.

Motion correction of the images shown in Fig 3 was performed using motioncor2 ^60^, CTF estimation was done with GCTF ^61^. XY drift was measured by calculating the Euclidean distance between frame shifts as outputted from motioncor2. The distances were summed up for each exposure count. Using ten micrographs, the total drift for each exposure count was used to calculate the mean value and standard deviation. Z change (using defocus change) was measured by calculating the defocus of a rolling average of aligned and summed frames. The number of frames to use was determined by testing different rolling averages for each sample type. For AmC images, a rolling average of 5 frames with a exposure of 4e-/Å^2^ was used, for GO images a rolling average of 7 frame with a exposure of 7e-/Å^2^ was used, and for OH images a rolling average of 9 frame with a exposure of 7.2e-/Å^2^ was use. The defocus from the first frame set was subtracted from all frames sets to establish the same start point for all images. Using ten micrographs, each corresponding frame set was used to calculate the mean value and standard deviation.

Eythrocruorin data (Fig 4, Fig 3 – figure supplement 1) were motion-corrected with motioncor2 ^60^, CTF estimation was done with GCTF ^61^, and particles were picked using Gautomatch (version 0.53, from K. Zhang, MRC-LMB, Cambridge). Processing was done in Relion3.0 ^62^. Initially, motion correction was performed with 5×5 patch correction, and CTF estimation was done on the whole micrograph. The initial set of 113,792 particles was subjected to 2D classification, and 46591 particles were selected for the initial model generated with C1 symmetry. This initial model was used as initial reference for refinement with D6 symmetry, which went to 4.9Å resolution. Per-particle CTF and astigmatism values were refined, and beam tilt correction was measured and corrected for, resulting in an improved resolution of 3.2Å ^62^. The particles were then subjected to 3D classification without alignment, from which a single class was selected containing 30,254 particles. Refinement of this subset of particles led to a resolution of to 3Å that improved to 2.9Å after polishing ^63^. 80S ribosome, apoferritin and TAF2 data was motion corrected using motioncor2^60^, CTF estimation was done with GCTF ^61^, and particles were picked using Relion LoG picker ^62^. Processing was done in Relion3.0 ^62^. Initially, motion correction was performed with 1×1 patch correction, and CTF estimation was done on the whole micrograph. Particles were subjected to 2D and 3D classification to select the best particles, followed by initial model generation, 3D refinement, CTF refinement, polishing ^63^, and alignment-free 3D classification. The final set of apoferritin particles was refined to 2.6Å and the final set of TAF2 particles was refined to 2.8Å. 80S ribosome refinement was subjected to multibody refinement ^64^ with to locally refine the 60S (2.7Å), 40S body (2.9Å) and 40S head (3.0Å). All local resolution calculations were done using Relion ^62^.

## Acknowledgements

Most of the data was collected at the Cal-Cryo facility. We thank Robert Louder for providing *Eisenia fetida* erythrocruorin; Patricia Grob, Robert Louder, Ben LaFrance, Vignesh Kasinath, and Basil Greber for discussion; Patricia Grob for electron microscopy support; Abhiram Chintangal and Paul Tobias for computing support; Patricia Grob and Giho Park for feedback on GO prep method; Robert Louder, Basil Greber, Vignesh Kasinath and Yuan He for comments on the manuscript. Images of cryo-EM reconstructions were generated using UCSF ChimeraX ^65,66^ (developed by UCSF RBVI with support from NIH R01-GM129325, OCICB and NIAID).

## Author Contributions

A.L. purified apoferritin and A.B.P purified TAF2. A.B.P devised a modified GO coating protocol and prepared, collected, processed, and analyzed cryo-EM data. A.B.P. D.T. assisted in tomogram collection. A.B.P wrote the paper with input from other authors.

---

**Figure 1 – Figure Supplement 1.**
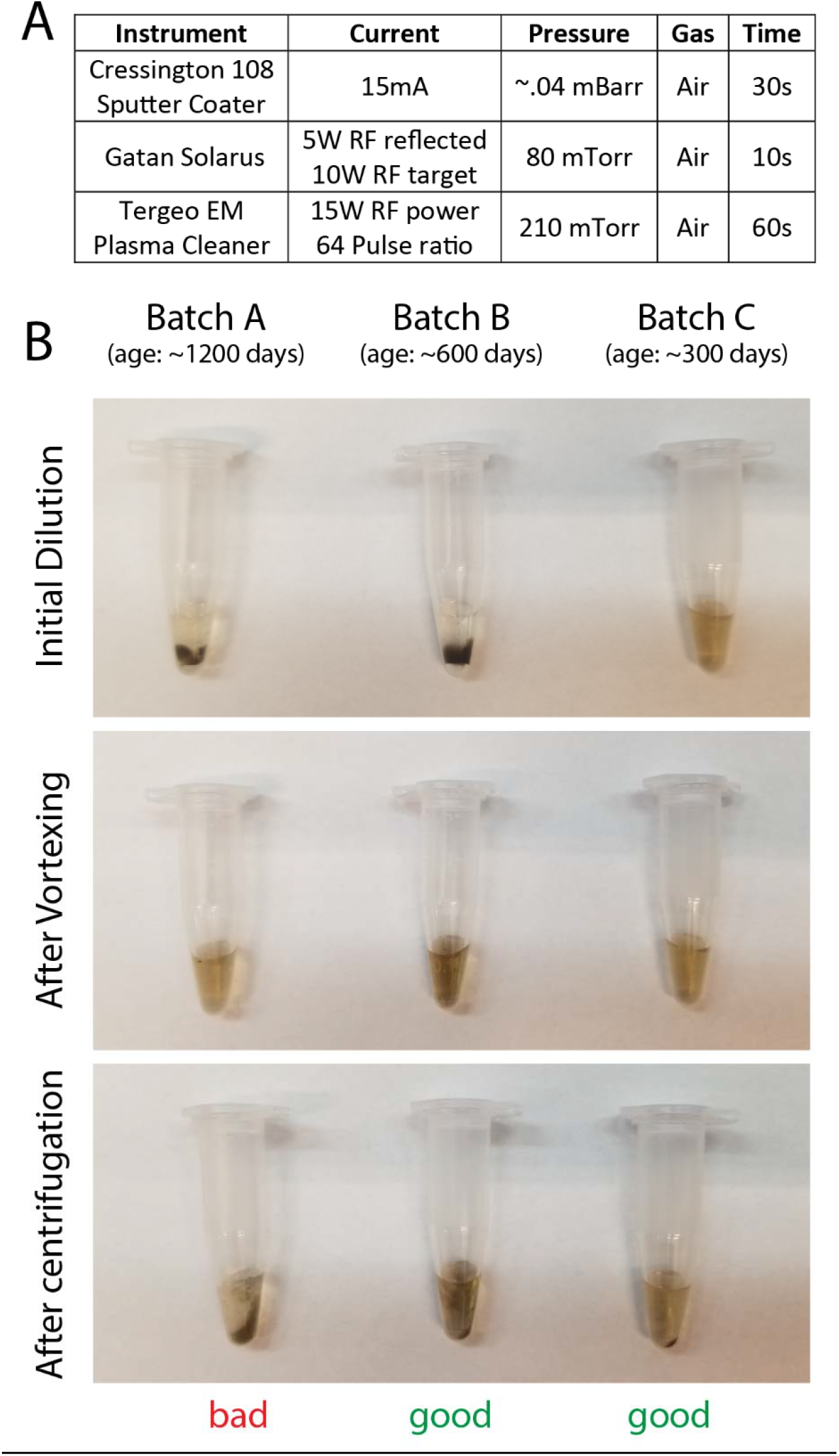
Grid preparation settings. A. Recommended glow discharge settings B. Graphene oxide dispersions batch and age variations (2mg/mL Sigma-Aldrich 763705-25ML)

**Figure 2 – Figure Supplement 1.**
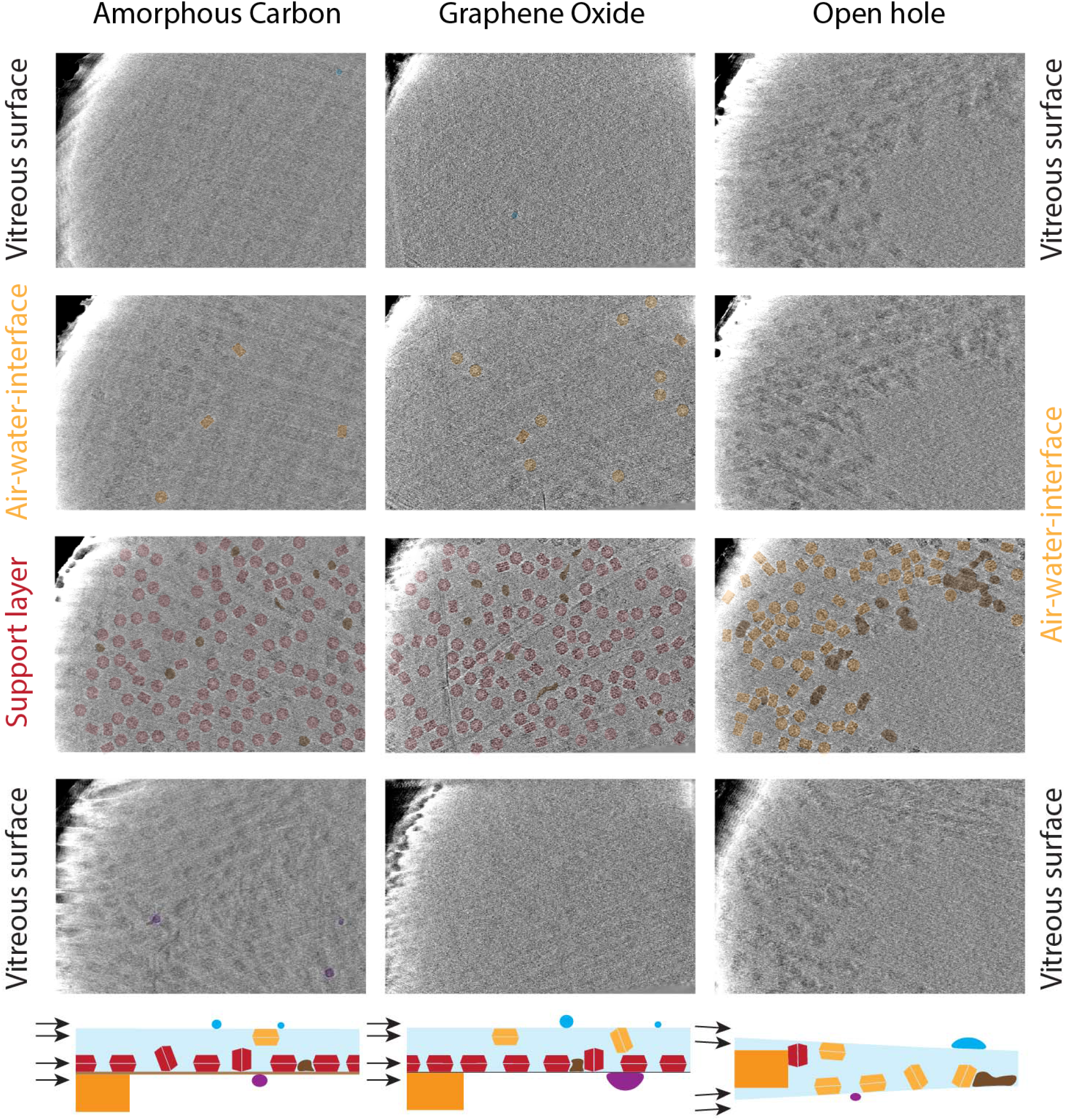
Tomographic slices at different heights in the cryo-EM sample. Tomographic slices from: (top row) above the vitreous ice layer; (second row) the top of the vitreous ice layer, showing particles bound at the air-water-interface on the blotted side; (third row) the bottom of the vitreous layer, showing particles either bound to the support layer (for AmC or GO grids) or to the air-water-interface (for OH grids) away from the blotted side; (bottom row) below the vitreous ice layer. Particles are colored based on their distance from the lower surface: red - particles bound to the support layer, orange - particles bound to air-water-interface, blue/purple – crystalline ice on the top/bottom of the vitreous ice layer, brown – damaged and aggregated complexes.

**Figure 2 – Figure Supplement 2.**
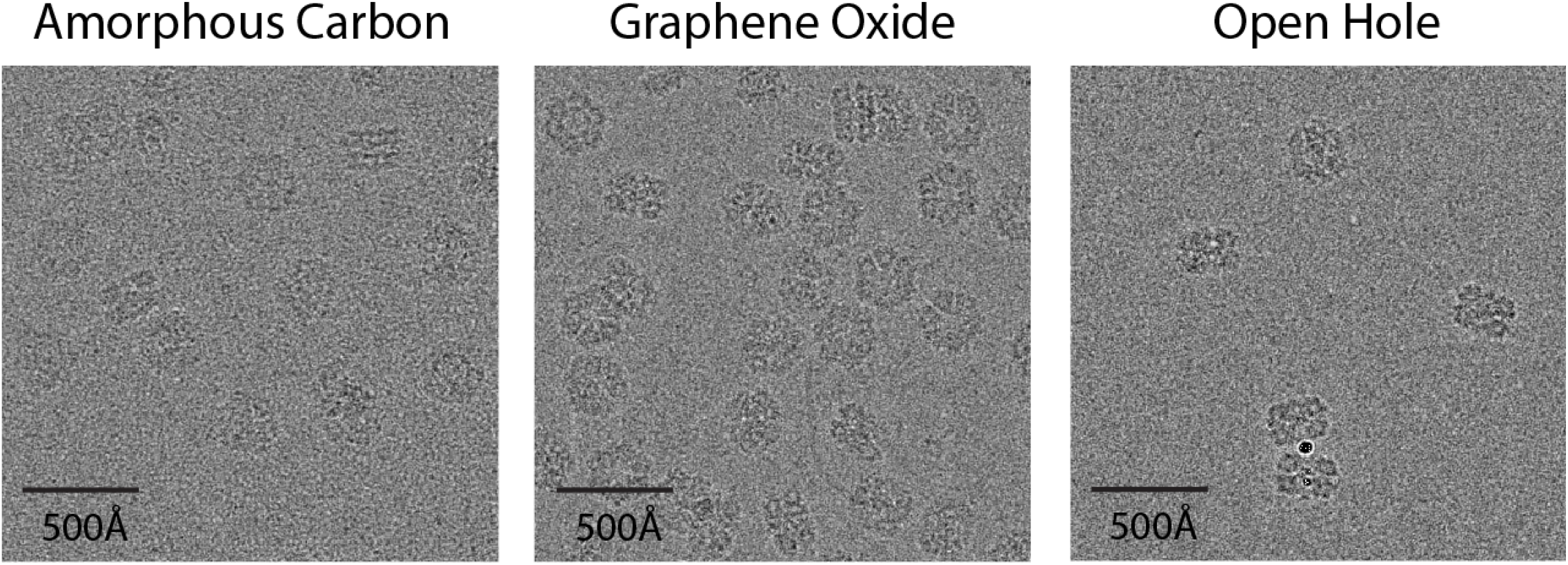
Support layer background. Representative micrographs from AmC, GO and OH grids. All images are at ∼1µm defocus.

**Figure 3 – Figure Supplement 1.**
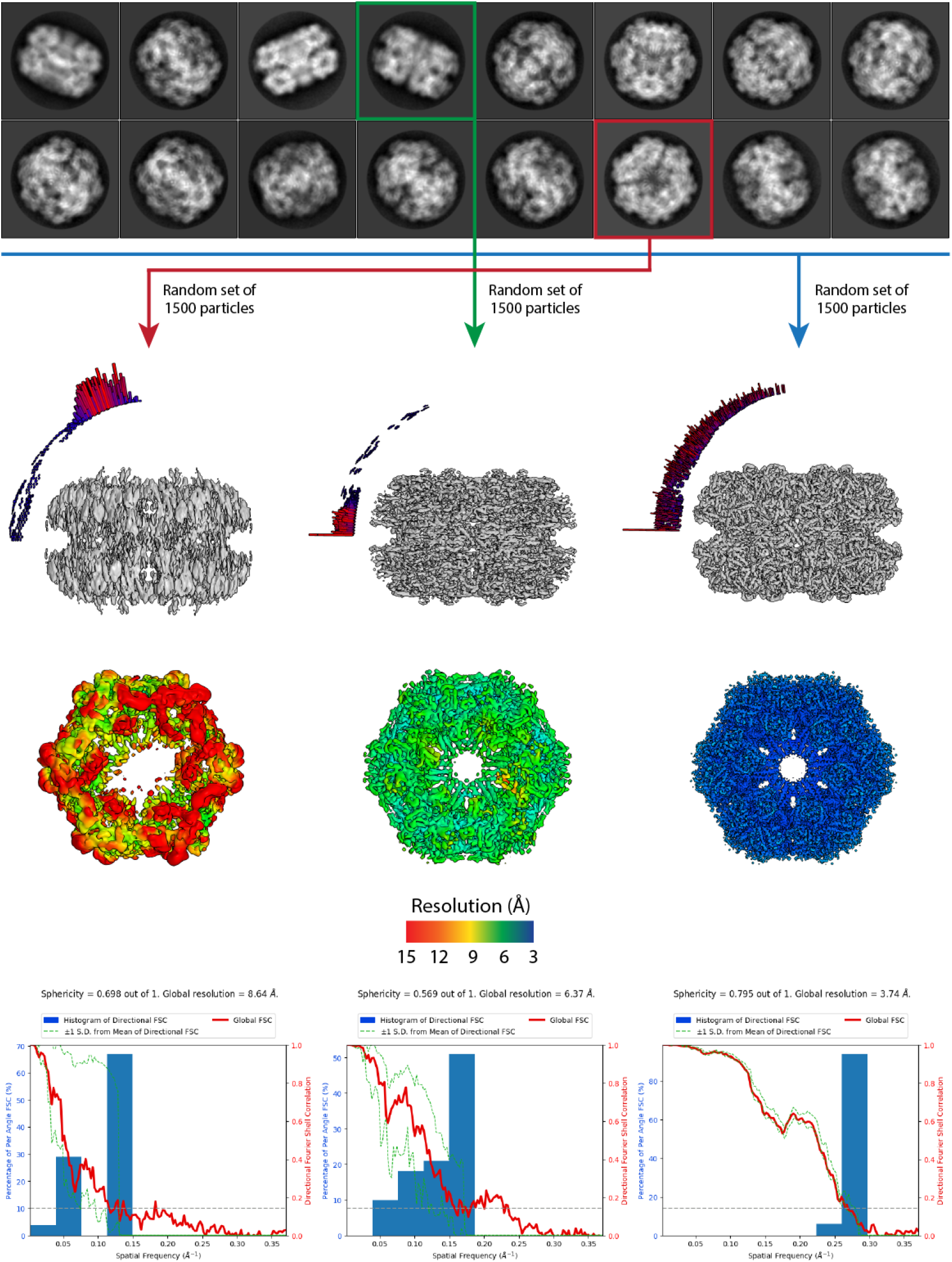
Effect of preferential views on 3D reconstructions. Erythrocrurin prepared on a graphene oxide coated UltrAuFoil grid collected at a 40° tilt. From select 2D class averages 1500 particles were used to generate a 3D reconstruction (with D6 symmetry). The reconstruction on the left was generated from a single 2D class average corresponding to the top view (down the C6 symmetry axis) resulting in an 8.6A reconstruction with a sphericity of .698 as measured by 3DFSC ^39^. The map from this reconstruction is very streaky with little interpretable secondary structure. The reconstruction in the middle was generated from a single 2D class average, perpendicular to the C6 axis, down the C2 symmetry axis, and thus containing six views around the particles C6 axis), resulting in a 6.4A reconstruction with a sphericity of .569. The map from this reconstruction had clearly visibly α-helices albeit with some features elongated. The right reconstruction contains a near complete set of views and resulted in map at 3.7A with a sphericity of .795 with clearly visible secondary structure (both α-helices and β-strands), and clear density for large side chains. Reconstructions colored by local resolution as calculated in Relion ^62^.

**Figure 4 – Figure Supplement 1.**
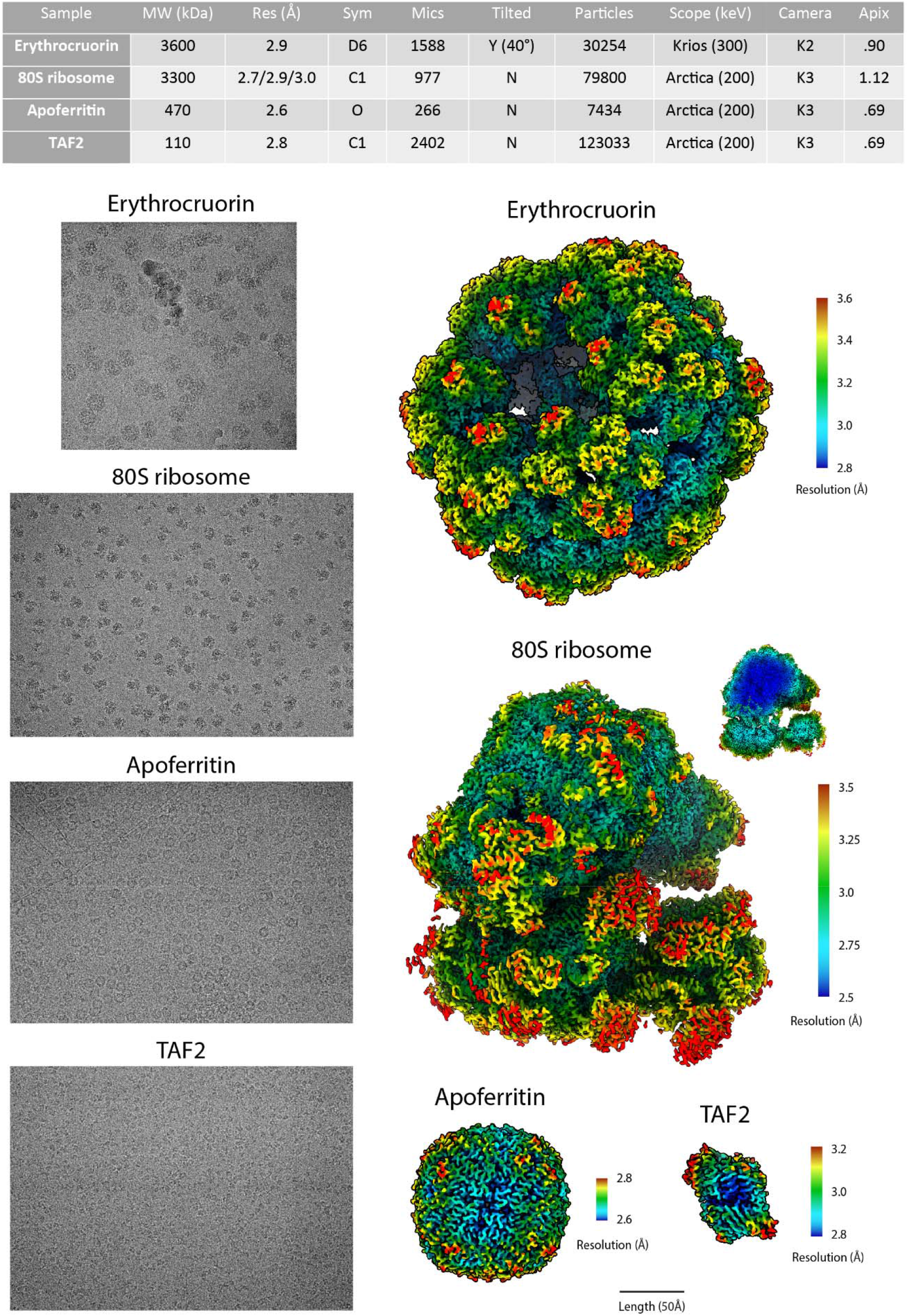
Reconstructions for different samples prepared on GO coated grids. Top: table summarizing the sample datasets collected on GO coated grids. Bottom left: example micrographs (all at ∼1.3µm defocus). Bottom right: reconstructions colored by local resolution as calculated in Relion ^62^. Structures shown to scale (except 80S ribosome cut through), scale bar at bottom.

